# Auditory efferents facilitate intensity change detection in humans: evidence from concurrent ear, brain, and behavior measures

**DOI:** 10.1101/2025.03.06.641896

**Authors:** Yuan He, Miranda J. Adamczak, Vijayalakshmi Easwar, Sriram Boothalingam

## Abstract

Feedback mechanisms in the brain, such as the medial olivocochlear reflex (MOCR), can aid signal detection and discrimination in noise in animals. Extrapolating these findings in elementary perception to speech understanding in noise abilities in humans has been inconclusive. To investigate the functional role of the MOCR in elementary auditory perception in humans, we concurrently measured ear (otoacoustic emissions) and brain (N1-P2 acoustic change complex [ACC]) responses while participants performed a behavioral intensity change detection task. The intensity change was imposed on a triad of ‘deviant’ clicks that occurred at early-, middle-, or late-temporal positions of a click train commensurate with null-, partial-, and full activation of the MOCR, respectively. We found faster behavioral detection and shorter ACC latencies for the middle- and late-occurring deviants relative to the early deviant. The improved behavioral detection of later-occurring deviants correlated significantly with the degree of MOCR activation and ACC latency. These mutual correlations suggest a direct involvement of the MOCR in human signal detection, underscored by earlier cortical detection. These novel findings advance our understanding of the mechanisms involved in speech perception in noise and aid in developing hearing assistive technologies.

## 1. Introduction

Feedback networks in the brain serve as a conduit for the central systems to influence bottom-up neural encoding. One such system, the medial olivocochlear reflex (MOCR), at the level of the brainstem, has been shown to be particularly relevant for auditory perception in animal models. Specifically, the MOCR improves neural encoding of, or unmasks, salient signals in background noise (reviews: Guinan, 2014a, 2018; Lopez-Poveda, 2018; Lauer et al., 2021). This putative unmasking function has been extrapolated to improve speech perception in noise in humans, albeit with equivocal findings (e.g., Kumar and Vanaja, 2004; Maruthy et al., 2017 vs. Mertes and Johnson, 2020; Rao et al., 2020). Given the redundancies in the neural encoding of speech, its perception is robust to internal (e.g., talker variation, pitch) and external (e.g., noise, reverberation) factors. As such, it is not surprising that a direct relationship between the MOCR activity, which is thought to improve peripheral auditory encoding in noise, and speech perception in noise is equivocal. To this end, here we ask whether the MOCR activation benefits more elementary auditory perception — intensity change detection — that, in turn, is known to contribute to speech perception.

The hypothesized role of the MOCR in elementary auditory perception and speech perception in noise stems from its ability to inhibit cochlear activity. This inhibition results in an improvement in the dynamic range of the auditory nerve, which is otherwise saturated by background noise (Winslow and Sachs, 1987; Liberman and Guinan, 1998). Unequivocal evidence for the contributions of this MOCR mechanism to hearing comes from direct electrical stimulation (Winslow and Sachs, 1987, 1988; Kawase and Liberman, 1993; Kawase et al., 1993; Delano et al., 2007; Seluakumaran et al., 2008) and lesion (Capps and Ades, 1968; Dewson, 1968; Hienz et al., 1998; Clause et al., 2017) studies.

Evidence from both animal and human studies has revealed a possible link between MOCR activation and enhanced elementary auditory perception. In animal models, activation of the MOCR has been shown to improve tone detection, intensity discrimination, frequency discrimination, and vowel discrimination in noise (Geisler, 1974; Winslow and Sachs, 1987, 1988; Dolan and Nuttall, 1988; Kawase et al., 1993, 1993; Micheyl et al., 1997; Hienz et al., 1998; Seluakumaran et al., 2008) A well-documented perceptual phenomenon in human psychophysics where signal detection is influenced by temporal positioning of target and masker, typically termed “overshoot”(Elliott, 1965; Zwicker, 1965) or “the temporal effect of masking” (Hicks and Bacon, 1992), has been postulated, at least in part, to result from the MOCR inhibition of cochlear activity (e.g., Bacon and Moore, 1986; McFadden and Champlin, 1990; Hicks and Bacon, 1992; von Klitzing and Kohlrausch, 1994; Strickland, 2001, 2004, 2008; Jennings et al., 2011; Lopez-Poveda et al., 2013; Roverud and Strickland, 2014; Marrufo-Pérez et al., 2018). More specifically, the signal detection in maskers is improved when either the masker precedes the signal in time, or an additional sound, called precursor, precedes the masker and signal combination. Neural adaptation (Bacon and Healy, 2000) and the MOCR mediation on adaptation (Strickland, 2004; Walsh et al., 2010a; Jennings et al., 2011) have been proposed as potential mechanisms that underpin the psychophysical overshoot. Auditory enhancement, where the improved signal detection in noise is due to the pre-cueing of spectral elements of the masker, is another phenomenon that may be related to the MOCR unmasking (Viemeister, 1980; Summerfield and Assmann, 1987; Thibodeau, 1996). Evidence of an MOCR involvement in the overshoot or enhancement effects comes from (1) the similarity in the time course of the MOCR (Liberman et al., 1996; Backus and Guinan, 2006; Boothalingam et al., 2021) and the temporal effects and (2) studies that measure both otoacoustic emissions (OAEs) and behavioral detection or discrimination in noise in the same group of listeners (overshoot: Keefe et al., 2009; Walsh et al., 2010; Henin and Long, 2013; enhancement: Beim et al., 2015). However, relationships between MOCR strength and perceptual effects are not consistently observed across studies [reported by Walsh et al. (2010) but not Keefe et al. (2009) and Beim et al. (2015)]. The discrepancy could be primarily due to the inadequate sensitivity of OAE measures when using stimuli designed for psychophysical overshoot conditions. A designated MOCR assessment method (e.g., with a contralateral noise elicitor) would be preferred (Henin and Long, 2013).

The facilitative MOCR influence on elementary auditory perceptual tasks in animals has been extrapolated to speech perception in noise for humans. Investigations on the relationship between MOCR activation and speech perception in noise in humans have produced mixed results. A positive correlation between MOCR activation and speech perception has been reported in some studies (Kumar and Vanaja, 2004; Maruthy et al., 2017) but not in others (Mertes and Johnson, 2020; Rao et al., 2020). The choice of speech materials and perceptual tasks can affect the observed MOCR involvement in speech perception (Hernández-Pérez et al., 2021). Prior psychophysical work in humans, which employed simple, nonsense stimuli and elementary perceptual tasks such as signal detection or discrimination, also suggests a potential MOCR role (Micheyl et al., 1995; Micheyl and Collet, 1996; Morand-Villeneuve et al., 2002; Keefe et al., 2009; Roverud and Strickland, 2010; Jennings and Strickland, 2012; Mertes et al., 2018, 2019; Wojtczak et al., 2019; Mertes and Johnson, 2020). However, behavioral results are not always paired with a direct measure of the MOCR or brain activity.

Several reasons may explain the discrepancy in the human data relating MOCR to perception. First, the assessment of the MOCR in humans typically relies on the non-invasive estimates of cochlear function, such as OAEs. In clinically normal-hearing ears, the observable changes in OAEs used to index the MOCR are small, typically 1 to 2 dB (Guinan, 2006). There is also no standardized approach for OAE-based estimation of the MOCR as studies employ various stimuli and paradigms. Second, MOCR strength and speech perception in noise are usually measured sequentially, with a myriad of speech materials across studies. This approach has recently been criticized (Guinan, 2014b) because MOCR activity can vary with task difficulty (Delano et al., 2007) and the type of speech task (Hernández-Pérez et al., 2021). Third, and perhaps most importantly, speech perception involves complex higher-level processing. Centers rostral to the MOCR may engage compensatory mechanisms even if the MOCR function is suboptimal (Hernández-Pérez et al., 2021; Boothalingam et al., 2023). This will potentially obscure any relationship between MOCR function and speech perception. As such, it remains unclear if the MOCR provides any direct perceptual benefit that may be related to speech perception in noise for humans.

To better understand the perceptual improvements that the MOCR may bring to humans, we investigated the role of MOCR on an elementary psychophysical ability essential for speech perception in noise — detecting intensity changes in time-varying signals. Temporal cues, such as envelope and fine structure cues carried in the transient intensity changes of ongoing signals, are important for robust and faithful neural speech encoding (Shannon et al., 1995; Turner et al., 1995; Xu and Zheng, 2007; Swaminathan and Heinz, 2012). Borrowing from psychoacoustics literature, we designed the current paradigm to study the influence of the MOCR on intensity increment detection (de Boer, 1986; Grose and Hall, 1997; Moore and Peters, 1997; Oxenham, 1997; Heinz and Formby, 1999). Further, to mitigate the aforementioned concerns in study design, we concurrently estimated (1) MOCR activity elicited by click trains and (2) cortical auditory evoked potentials, i.e., the N1-P2 acoustic change complex (ACC), evoked by intensity change embedded in a click train, while participants also behaviorally detected the transient intensity change. We measured the ACC concurrently for two reasons: (1) to assess the neural processing underlying behavioral detection of intensity changes within click trains, and (2) to investigate whether MOCR strength would predict cortical, in addition to behavioral, detection. We hypothesized that the MOCR facilitates behavioral detection and cortical registration of intensity change.

## 2. Material and Methods

### 2.1 Participants

A total of 25 young adult (age: 23.2 ± 2.3 years; 22 females, 3 males) participants with clinically normal hearing completed the study in three sessions, 1.5 - 2 hours each. All participants had normal hearing as defined by hearing thresholds (< 20 dB HL between 0.25 - 8 kHz; AD629 diagnostic audiometer, Interacoustics, Denmark) and normal middle ear function (tympanogram peak pressure: ± 100 daPa and static admittance between 0.2 - 1.6 ml; Titan, Interacoustics, Denmark). All participants were also required to have measurable distortion product (DP) OAEs (1 - 8 kHz for 55/65 dB SPL stimuli; SmartDPOAE, IHS, FL, USA). All study procedures were approved by the Health Sciences Institutional Review Board at the University of Wisconsin-Madison. All participants provided written informed consent prior to participating in the study.

### 2.2 Stimuli

Participants were presented bilaterally with a train of clicks in a task of intensity change detection while their OAEs and EEG evoked by the same stimulus were measured concurrently. Clicks were band-limited (0.8 - 4.8 kHz) and presented at 62.5 Hz continuously for 1.536 s (Boothalingam et al., 2021). The click rate equates to an ‘epoch’ duration of 16 ms, i.e., the duration between the onsets of successive click presentations. This specific rate was used to present sufficient clicks to activate the MOCR while also sampling OAEs in each epoch (Boothalingam et al., 2021).

The level of click trains was calibrated in each participant’s ears using the forward pressure level (FPL) method (Scheperle et al., 2008; Rasetshwane and Neely, 2011; Boothalingam and Goodman, 2021). FPL was used to shape the click within the 0.8-4.8 kHz band such that it was spectrally flat at the tympanic membrane in each participant. This procedure was done to minimize between-subject variability arising from the ear canal and middle ear acoustics (Boothalingam and Goodman, 2021; Boothalingam et al., 2022). Clicks were digitally generated in MATLAB (Mathworks, MA, USA) on an iMac, converted to analog signals (at 96 kHz; Fireface UFX+, RME, Germany), and delivered to participants’ ears through an ER10X probe system (Etymotic Research, IL, USA). Bilateral ER10X probes were securely placed in participants’ ears by (1) suspending them from the ceiling of the sound booth, (2) attaching them to hollowed-out earmuffs (Mpow, CA, USA), and (3) puttying around the in-ear probes (Silicast, Westone Laboratories, CO, USA). The ER10X probe microphones registered and pre-amplified (+20 dB) the ear canal acoustic pressure before feeding it to the soundcard (at 96 kHz; Fireface UFX+, RME, Germany) for analog-to-digital conversion and storing in a secure hard disk for offline analyses. Ear canal pressure contained both the stimulus and the evoked OAEs.

Cosine-squared onset and offset ramps were imposed on the first and last 6 clicks, i.e., 96 ms, to minimize onset and offset cortical responses (Onishi and Davis, 1968; Skinner and Jones, 1968). In the detection task, two types of click trains were presented: (1) standard click trains, which contained equal-level clicks presented at 75 dB peak-to-peak (pp) SPL, and (2) deviant click trains, which were characterized by the presence of a triad of clicks with an intensity increment embedded in the standard click train. The deviant click triad occurred at one of the three temporal locations in a given trial – early (96 ms), middle (528 ms), and late (1008 ms) – coinciding with null-, part-, and full-activation of the MOCR, respectively (Fig 1E). Figs 1B-D. illustrate the stimulus used in the study and the three temporal positions of deviant click triads. The stimulus was intentionally designed to continue after the late position to avoid offset responses in the EEG contaminating the ACC at the late position (Onishi and Davis, 1968). A 1.5-second inter-stimulus interval (ISI) followed each click train to allow a complete reset of the MOCR (Backus and Guinan, 2006; Boothalingam et al., 2021) and for neural/cortical activity to return to baseline.

**Fig 1.**
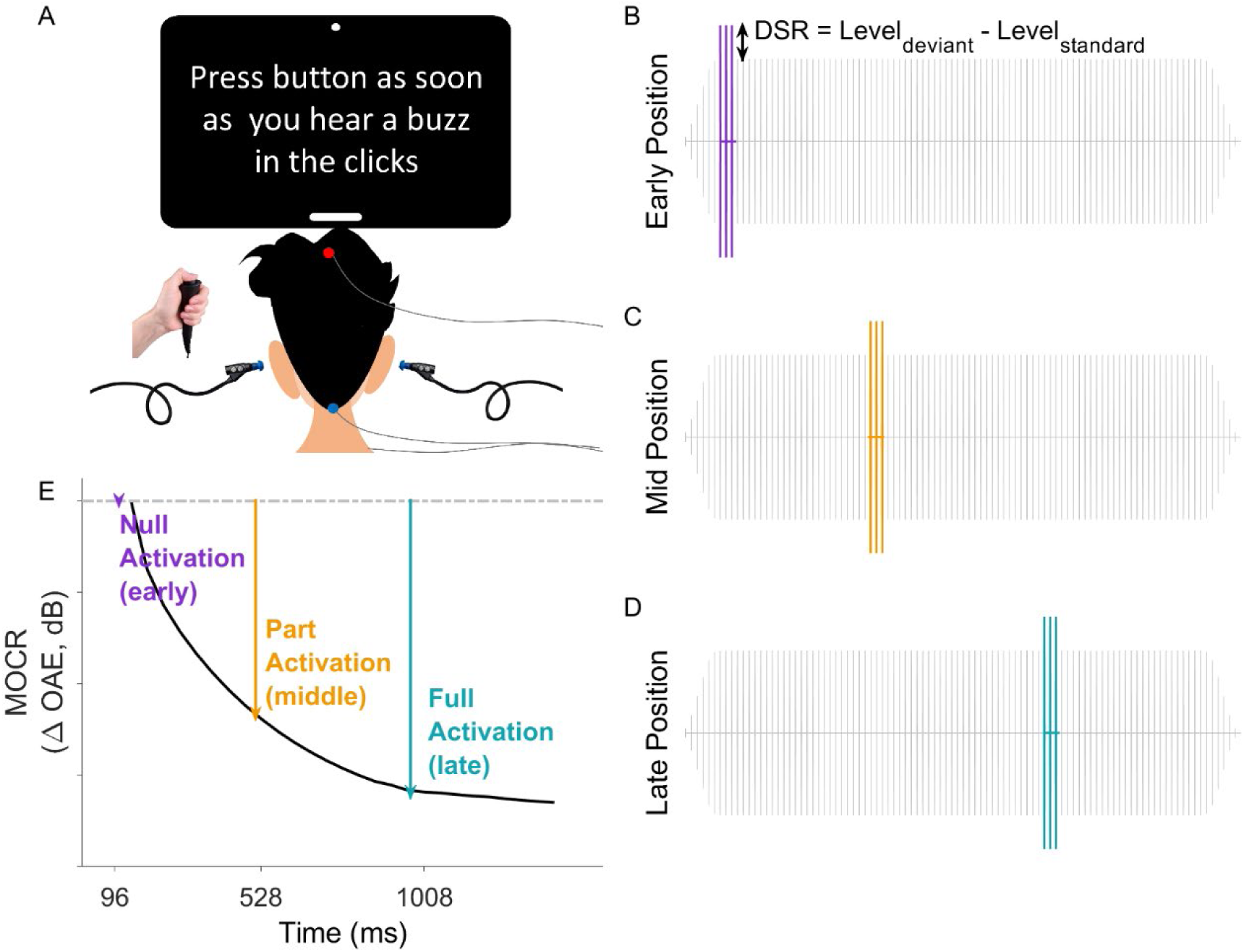
Illustration of experimental design (A) and stimuli (B-D). Listeners were to detect deviant clicks embedded in a click train and indicate detection by pressing a response button. In Phase 2, bilateral otoacoustic emissions were acquired by inserting probe earphones, and, concurrently, EEGs were recorded from the Cz (non-inverting) and nape (inverting) positions. Panels B-D are examples of click trains with deviant click triads embedded in different temporal positions of a standard click train: early position: 96 ms, middle position: 528 ms, and late position, 1008 ms. DSR is calculated by subtracting the peak RMS amplitude of the standard expressed in dB SPL from that of the deviant expressed in dB SPL. In panel E, the expected MOCR activation, indexed using OAEs, is schematized. Note that data obtained for each position is color-coded in all later figures: purple for early-positioned deviants, orange for middle-positioned deviants, and cyan for late-positioned deviants.

Evoked EEG was concurrently acquired with a wide passband (1-5000 Hz) and amplified (10^5 times) by the OptiAmp system (Intelligent Hearing Systems [IHS], FL, USA) before feeding the analog signals to the soundcard for digitization (at 96 kHz; Fireface UFX+, RME, Germany). Three sintered Ag-AgCI electrodes with a vertical montage were used for EEG acquisition. The vertex (Cz) served as the non-inverting site for the inverting site at the posterior midline of the neck (nape), and the left collarbone served as the ground site (Fig 1A). Before the start of test sessions, the electrode sites were cleaned with alcohol wipes and a mild abrasive gel (Nuprep; Weaver & Company, Aurora, CO) to lower the scalp-electrode impedance. Electrode impedances were monitored throughout the test session and were always below 3 kΩ across all electrodes.

### 2.3 Procedure

The study consisted of three phases: (1) familiarization, (2) threshold estimation, and (3) main experiment. In all phases, the listeners’ task was to detect the intensity change that occurred in any of the three pre-selected positions of a click train (Figs 1B-D). Participants were instructed to press a button as quickly and accurately as possible whenever they thought they heard a buzz associated with the occurrence of a brief intensity change, i.e., the deviant (Fig 1A). Therefore, a single interval, Yes/No task was used to measure the deviant detection in all study phases.

In the familiarization phase (Phase 1), listeners were briefly trained to distinguish between the standard and deviant stimuli using sound demos. The intensity of the deviant click triads was kept at a fixed level (10 dB higher than the standard clicks) in the sound demos so that the intensity change was obvious to naive listeners. This provided the opportunity for the participants to familiarize themselves with the acoustics of the deviants in the ongoing click train. The intensity level difference between the deviant and standard clicks is henceforth referred to as the deviant-to-standard ratio (DSR). To graduate to the next phase, all listeners were then asked to complete a detection task and achieve at least 80% accuracy. The detection task in Phase 1 contained 20 click trains, with 50% being deviants. All participants successfully graduated to the subsequent study phase. In the threshold estimation phase (Phase 2), individualized detection thresholds, defined as the minimum detectable DSR, for deviants at all three temporal positions were measured using the method of constant stimuli. For each temporal position, the detection accuracy was measured for deviants with a DSR between 1 and 7 dB in 1 dB steps. The range of DSR was chosen based on pilot work to avoid floor and ceiling effects. There was a total of 140 click trains (20 trains/DSR × 7 DSRs) with a 50% probability of the deviant occurring for each temporal position. The order of deviant trains was randomized across test conditions and participants. A psychometric function was then constructed for each temporal position and participant to estimate the detection thresholds (Wichmann and Hill, 2001a).

In the main experiment phase (Phase 3), the accuracy and reaction times (RTs) of detecting deviants, MOCR activity, and ACC in response to deviants, were measured concurrently at a suprathreshold level. Ideally, the detection sensitivity (d’) would be equalized across three temporal positions to reduce the effect of task difficulty on the measurement of RT and MOCR (McGarrigle et al., 2014; Alhanbali et al., 2019; Hernández-Pérez et al., 2021). We measured the deviant detection in equal-SL and equal-SPL conditions and hypothesized that the task would be more equalized in the equal-SL condition. In the equal-SL conditions, the deviants were presented at 6 dB SL (6 dB above the individualized detection threshold estimated in Phase 2) for each temporal position. In the equal-SPL condition, the deviant levels were the same for all temporal positions and determined by the highest detection threshold among all three temporal positions for each individual. For example, if a participant’s thresholds were 4, 3, 2 dB DSR for the early, middle, and late positions, respectively, the presentation levels for the deviants in the equal-SL condition were 10, 9, 8 dB DSR for three positions, and 4, 4, 4 dB DSR if in the equal-SPL condition. Note that the data from the equal-SPL condition (not reported in Figs 3-5) was only used to confirm our hypothesis that detection sensitivity and overall task difficulty in Phase 3 would be equalized across temporal positions in the equal-SL condition. Therefore, any perceived position effects in the concurrent measures will be more likely to arise from a true MOCR influence and less confounded by perceptual sensitivity and task difficulty.

The detection of three different deviants were measured in a block design. For each temporal position, a total of 400 click trains with 20% deviant probability were presented in two blocks, yielding 80 trials for each deviant. Table 1 summarizes the different paradigms in three study phases. An oddball task of detecting the deviants was used to reduce the anticipation effect and optimally evoke the ACC responses to each temporal position. Here, oddball refers to the participants’ task in detecting the randomly occurring click trains with deviants (20%) among trains with no deviants.

**Table 1.** Summary of the experiment paradigm in three study phases. The listening task in all phases was the same. Each temporal position was measured separately in a block design.

| Study Phase |  | Listening Task | Deviant-to-standard ratio<br>(DSR) | Deviant<br>occurrence<br>rate (%) | Total Trial<br>Number |
| --- | --- | --- | --- | --- | --- |
| 1 | Familiarization | Press a<br>response button<br>when<br>participants<br>detected the<br>intensity change<br>in the click train,<br>as quickly and<br>accurately as<br>possible | 10 dB | 50 | 20 |
| 2 | Threshold<br>Estimation |  | 1 to 7 dB in 1 dB<br>steps | 50 | 140 (20<br>per DSR<br>level) |
| 3 | Main<br>Experiment |  | Detection<br>Threshold (DSR)<br>at each position +<br>6 dB (equal-SL) | 20 | 400 |

Throughout the test sessions, participants were seated in a sound-attenuated booth and were given visual feedback on a monitor in the form of performance scores. Participants were also instructed to minimize unnecessary head and body movements and stay relaxed during the test. Adequate breaks were provided upon request to avoid fatigue.

### 2.4 Data analysis

#### 2.4.1 Behavioral intensity change detection

A four-parameter cumulative Gaussian distribution was fitted to individual data to estimate the underlying psychometric functions. We used the model described by (Wichmann and Hill, 2001a),

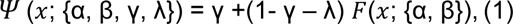

where *x* denotes the DSR (dB), and the four parameters *α, β, γ*, and *λ* determine the overall shape of the fitted psychometric function. The function F here was chosen to be the cumulative Gaussian distribution. Hence, *α* and *β* describe the mean and standard deviation of the distribution. Parameters *γ* and 1-λ describe the lower and upper asymptotes of the psychometric function. The lower asymptote *γ* corresponds to the false alarm rate in the currently used Yes/No paradigm. In the fitting process, λ was set to a small value, 0.1, to minimize slope bias. Parameters of our main interests were the detection thresholds 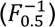 estimated at 50% accuracy and the standard deviation that roughly reflects the spread and the slope of the distribution. The goodness of fit was then assessed by Bayesian inference methods, implemented in the Psignifit (toolbox for Bayesian psychometric function estimation, version 4, MATLAB) (Wichmann and Hill, 2001b; Schütt et al., 2016).

In Phase 3, the elapsed time between the onset of a click train and the button press was obtained as the reaction time (RT). RT data were pooled across conditions for each participant to identify and reject outliers. Outliers were button presses that occurred earlier than 100 ms (Luce, 1986; Whelan, 2008) from the deviant or slower than 4 standard deviations away from the mean RT. An exponential (ex)-Gaussian distribution was fitted to individual RT data because it provides a more accurate characterization to the RT data compared to a single Gaussian distribution-(DISTRIB MATLAB toolbox; Lacouture and Cousineau, 2008). This distribution has a convoluted form of exponential and normal distributions and therefore has been recommended over standard central tendency estimators for describing multiple sequential cognitive processes (Luce, 1986; Lacouture and Cousineau, 2008; Matzke and Wagenmakers, 2009; Castro-Palacio et al., 2021). Three parameters *µ, σ*, and *τ* were estimated from each fitted distribution. The mean and deviation of the Gaussian component were denoted by *µ* and *σ* whereas the mean of the exponential component was denoted by *τ*. The goodness of fit was incorporated into the model fitting procedures. The search for best-fitting parameters would only terminate when the estimated goodness of fit reaches a predetermined criterion (tolerance = 0.0001; Lacouture and Cousineau, 2008).

In addition to the RT measures, detection accuracy was measured for the suprathreshold detection. D-prime (d’) was calculated for each position and corrected for the perfect accuracy cases (Hautus, 1995; Macmillan and Creelman, 2004). Extreme false alarms and hit rates (0 or 100%) were adjusted to avoid infinite d’ values.

#### 2.4.2 OAE and MOCR analysis

OAEs were analyzed to estimate the magnitude of MOCR activation using the methods described in (Boothalingam et al., 2021). Briefly, the measured pressure from the ear canal was bandpass filtered between 0.8 and 4.8 kHz, i.e., the same as the click stimulus. Clicks presented during the on- and off-ramp portions of the click train (96 ms × 2) were excluded from the OAE analysis due to their unequal stimulus levels. Therefore, time zero was redefined as the start of the first steady level click at 96 ms, and the total duration was reduced to 1.344 s. Additionally, the deviant click triad within a train was removed as these clicks are of a higher pressure level. The rest of the recorded pressure responses were then divided into click-length epochs of 16 ms. An interquartile range-based artifact rejection was employed, where epochs with root-mean-square (RMS) amplitude > 2.25 times the interquartile range were rejected. Less than 10% of the epochs were rejected using this approach in all position conditions and participants. Next, three consecutive epochs were averaged, i.e., decimated, to improve the signal-to-noise ratio (SNR), theoretically ∼4.77 dB, at the cost of MOCR time resolution, i.e., 48 ms from 16 ms. Decimation by three clicks were chosen as a compromise between reducing experiment time, obtaining at least 1000 click epochs at each time point along the time course to have high enough OAE SNR, and minimizing any effect on the time resolution of the MOCR time course. The reduction in time resolution is well within the onset time of the MOCR (Backus and Guinan, 2006; James et al., 2005), and as such our central assumption of capturing OAE without any MOCR activation (the time zero reference, explained below) would not be violated, resulting in a true representation of the MOCR magnitude at later time points along the time course.

OAE waveforms were extracted between 4.5 and 15 ms post click onset (peak of the click) in each averaged epoch and 1 ms on/off Hann ramps were applied to avoid spectral splatter. The resulting waveforms that had 12 dB SNR were considered in the time-frequency domain to extract only the expected OAEs based on average human cochlear delays (Shera et al., 2002, 2010) between 0.8 and 2 kHz where the MOCR effects are most pronounced (Lilaonitkul and Guinan, 2012; Zhao and Dhar, 2012). For each individual, OAEs were averaged in two dimensions. (1) within each epoch (4.5-15 ms) and (2) across conditions and click train repetitions (n=400 with deviant clicks removed). The within-epoch (RMS) average reduced the OAE into a single data point at the respective time along the click train. The averaging across conditions and click trains, while preserving the time course of the OAE change, i.e., MOCR, provided an improvement in overall SNR, similar to any OAE/EEG averaging. Although clicks were presented in both ears concurrently, we only report data on one ear per participant for computational simplicity as the MOCR and middle ear muscle reflex (MEMR) produce identical effects in the two ears using the time-course-based approach (Backus and Guinan, 2006; Boothalingam and Goodman, 2021; Boothalingam et al., 2021).The laterality of the chosen ear was counterbalanced between participants. Averaged OAE amplitude across time was referenced to the OAE at time zero, henceforth referred to as ΔOAE. That is, each data point in the ΔOAE was estimated by subtracting the OAE amplitude at time zero from the amplitude at every time point. Our assumption is that the MOCR is not activated at time zero, i.e., the first 48 ms. As such, referencing all subsequent OAE amplitudes to this first time point is likely to capture a true MOCR time course. If the MOCR were not activated, ΔOAE would be a straight line with a constant value of 0 across the duration of the click train. If the MOCR were activated, a reduction in amplitude, approximated by a two-term exponential fit can be expected (Liberman et al., 1996; Kim et al., 2001; Backus and Guinan, 2006; Boothalingam et al., 2021). Goodness of the fits were estimated by a resampling-based bootstrapped (x1000 times) implementation of the Heller-Heller-Gorfine (HHG) test. Briefly, this test assesses the hypothesis that the fit estimated data and the raw data come from the same distributions, which would be true if the fit approximates the data accurately. The p-values <0.007 (0.05/7) were considered as significant after applying Bonferroni corrections for performing multiple (7) comparisons, i.e., at each frequency band in a given individual. This process led to the rejection of two participants (n = 23 after rejection) who did not demonstrate significant MOCR at any of the 7 frequencies.

Finally, the amplitude of the MOCR at three temporal positions (early, middle, and late) was then extracted from the two-term exponential fit to index the MOCR strength. We extracted this data from the fit, rather than the raw data points, as means to minimize the inherent variability in ΔOAE. An individual analysis of all significant exponential fits and their corresponding residuals are presented in Supplementary Fig 1.

#### 2.4.3 MEMR analysis

The commonly used stimulus and MOCR elicitor in the OAE-based assays of MOCR measurement can often activate MEMR (Guinan, 2014b, 2018). As a result,, the observed MOCR effects are likely to be contaminated by MEMR activation. In addition, MEMR activation produces similar physiological changes, i.e., along the same time course and magnitude, on OAEs to that is produced by MOCR activation. Because separating MOCR and MEMR effects on OAEs is non-trivial, it is prudent to avoid MEMR activation using moderate-level stimuli that is less likely to activate the MEMR, as used in the present study, and check for potential contamination using sensitive methods (Boothalingam and Goodman, 2021). We applied the same time-course analysis to the click stimulus, instead of the OAEs, to estimate MEMR activation. The resulting MEMR strength is denoted as |Δstim|. Here we considered the full spectrum of the clicks as the MEMR affects all frequencies. The changes were also considered in absolute terms because the impedance changes caused by the MEMR can increase or decrease the stimulus level in the ear canal as a function of frequency. A |Δstim| of 0.12 dB is thought to be the threshold of MEMR potentially influencing MOCR estimates (Abdala et al., 2013). In Fig 3B, we plot the relationship between the MOCR (|ΔOAE|) and MEMR (|Δstim|) amplitude for the late position, where both reflexes were fully active. The relationship between the MOCR and the MEMR is non-significant both with and without the three participants (with: r = 0.34, p = 0.11; without: r = 0.36, p = 0.11), suggesting that the MEMR did not influence MOCR estimates. However, taking a conservative approach, we excluded these three participants for further analyses on the relationship among concurrent measures (n=20 after the second rejection). As such, the ear-brain-behavior link in detecting transient intensity changes reported in this work are predominantly driven by the MOCR and not the MEMR.

#### 2.4.4 ACC analysis

EEG data collected in intervals when participants successfully detected a deviant (a Hit response) were first downsampled to 1000 Hz. The entire EEG waveform was then segmented into epochs of 3.036 seconds (1.536-s stimulus duration+ 1.5-s ISI) and low-pass filtered at 15 Hz. An interquartile range-based artifact rejection, the same as that used for OAEs, was employed. EEG was then averaged across odd- and even-numbered epochs. ACC was aggregated from the average of all epochs. The noise floor was estimated from the difference between the averaged odd-numbered and even-numbered epochs.

A custom automated peak detection algorithm was used to roughly identify the ACC peaks. The algorithm would first localize the positive component (P2) within 300 ms of the onset of the deviant click triads and continue to search for a trough (N1) within −150 ms of the marked P2. The identified N1-P2 complex was then visually confirmed and re-picked, if necessary, by two independent observers (authors YH and VE). A third observer (author SB) resolved any disagreements. The N1-P2 peak-to-peak amplitude and latencies for each component were extracted for each participant and test condition.

#### 2.4.5 Statistical analysis

Statistical analyses were performed in R (R 4.2.1., Development Core Team 2022) using the linear and generalized linear mixed-effects model fitting packages (*lme4* package; Bates et al., 2015). To assess the effects of different temporal positions on all concurrent measures, we used a linear mixed model (LMM, ‘lmer’ function) with a repeated measures design (Laird and Ware, 1982). Participants were treated as a random effect in the model. Each measure was predicted by a fixed effect of deviant positions and a random effect of participants. Post-hoc Tukey test was conducted to compare pairs of deviant positions.

To assess the association among all concurrent measures, a repeated measures regression was performed using LMM models. All LMM models included a random effect of participants and allowed for random intercepts for participants as individualized “baseline” and random slopes for the linear association. Three separate models estimated the association between variables across three positions (1) RT and the OAE amplitude, (2) N1 or P2 latency and the mean RT, and (3) N1 or P2 latency and the OAE amplitude. Effect sizes were calculated for each fitted model using the ‘anova_stats’ function (as implemented in the *sjstats* package; Levine and Hullett, 2002; Tippey and Longnecker, 2016). Regression coefficients (*β*) with 95% CI are reported. Degrees of freedom in the fitted models were estimated by the Satterthwaite approximation approach (as implemented in the *lmerTest* package; Kuznetsova et al., 2017). Homoscedasicity, residual normality, and influential cases were examined for all fitted models.

## 3. Results

### 3.1 Behavioral intensity change detection thresholds

The detection threshold for the intensity increment at each temporal position was defined as the deviant-to-standard ratio (DSR), i.e., the minimum intensity change, from the standard click level, required to detect the presence of deviant clicks (illustrated in Figs 1B-D). Detection thresholds were estimated as the 50% correct point on a psychometric function fitted to individual data. The estimated individual thresholds and group trends are plotted in Fig 2A. A linear mixed model analysis revealed a significant effect of position (F [2, 43] = 15.5, p < 0.001) on the detection thresholds. The detection threshold for the early deviant position was the highest (worst) compared to that for the middle and late positions (p < 0.001 for both pairs as indicated by post hoc Tukey’s tests). The thresholds for the middle and late positions were not significantly different from each other (t (1, 46.1) = 0.4, p = 0.905). Detection thresholds (mean ± 1 SD) for the early, middle, and late positions were 4.2 ± 1.4, 3.5 ± 0.8, and 3.3 ± 0.9 dB, respectively.

**Fig 2.**
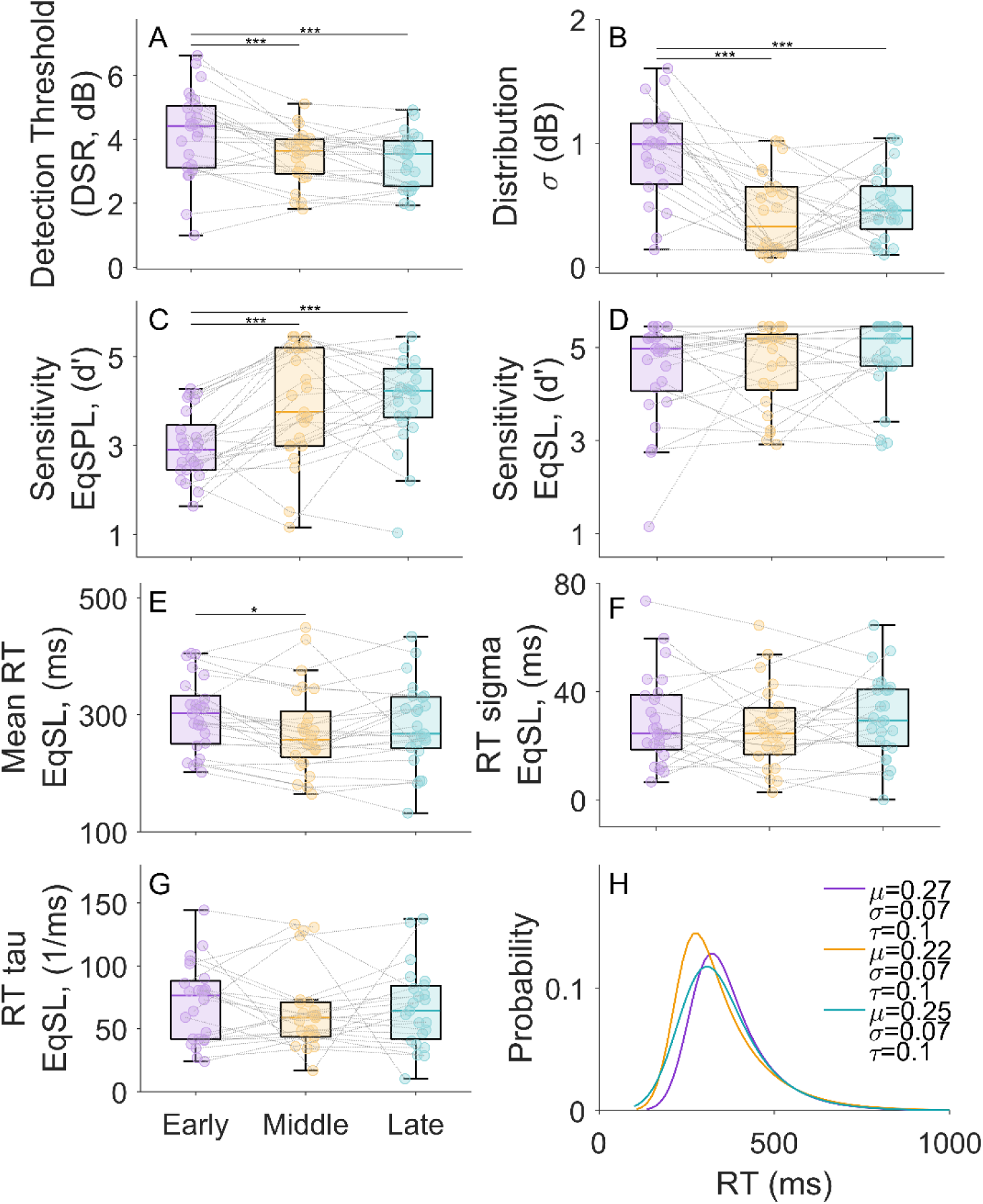
Behavioral detection of intensity change in click trains. Individual and group average (n = 23) detection thresholds obtained in Phase 2 for deviants at three temporal positions (early, middle, and late positions) are shown in panels A and B. Thresholds (A) were estimated from the 50% accuracy level on the psychometric functions. The spread (slope) of the fitted psychometric functions is shown in (B). (C) Detection sensitivity (d’) was measured for the equal SPL condition, where deviant levels were the same for all temporal positions and determined by the highest detection threshold of the three temporal positions. (D) d’ measured for the equal SL condition, in which deviant levels were 6 dB above the detection threshold for each temporal position. (E-G) Suprathreshold detection performance was assessed by the RT measures in the main experiment. Ex-Gaussian distributions were fitted to individual RT data and the mean, standard deviation (sigma), and exponential component (tau) of the distributions were plotted. (H) An example of a fitted individual RT distribution for one of the test conditions. \*\*\**p* <=0.0001, * *p*<0.05.

The standard deviations of the fitted psychometric functions (as shown in Fig 2B), were estimated by the differences between the DSRs measured at 75% and 25% correct points on the psychometric function. The estimated standard deviations reflect the spread of the psychometric functions and the overall steepness of the slope. A significant position effect (F [2, 43] = 18.0, p < 0.001) was observed between the early and the two later positions (post hoc Tukey’s test, p < 0.001 for both comparisons). No significant difference was found between the two later positions (t (1, 46.1) = −1.0, p = 0.606). The psychometric function for detecting the two later-positioned deviants was steeper than that for the early-position. This means that the rate of improvement in detectability of the early-positioned deviants with increasing DSR is lower relative to later-positioned deviants. The steepness of the psychometric function slopes has been demonstrated as a reflection of the cochlear mechanical nonlinearity (Schairer et al., 2003, 2008; Rodríguez et al., 2011). As indicated in the Schairer et al. (2003) model (demonstrated with forward-masking experiments), a low masked detection threshold or better detection performance associated with a steep psychometric function are results of the signal passing through the less compressive portion of the basilar membrane input-output function. If this is the case, the steeper psychometric function measured at the two later-positioned deviants may be attributed to the reduced peripheral compression (increased linearization) due to the MOCR-induced reduction of cochlear gain (Fletcher et al., 2016). Overall, the improved detection thresholds and steeper psychometric functions for the later-positioned deviants appear consistent with an MOCR involvement.

### 3.2 Suprathreshold intensity change detection

The difficulty of the detection task was equalized across individuals and temporal positions by presenting the deviants at equal SL, ensuring that intensity increments were easily and equally detectable at all three positions. This equalization was confirmed by comparing the detection sensitivity (d’) in an equal-SL condition with that in an equal-SPL condition. The d’ for the equal-SPL and equal-SL conditions are plotted in Figs 2C and 2D. The temporal position had a significant effect on the d’ in the equal-SPL condition (Fig 2C; F [2, 39] = 14.0, p < 0.001) but not in the equal-SL condition (Fig 2D) (F [2, 35] = 1.5, p = 0.231). This dichotomy indicates that the task difficulty was effectively equalized across temporal positions when the deviants were presented at equal SL. In the equal SPL condition, the later positions likely showed better sensitivity (re: the early position) because the highest DSR almost always occurred at the early position providing a level-related advantage for later positions. We thus mitigated threshold differences, potentially driven by difference in stimulus energy and salience, between temporal positions that may conflate a true MOCR position effect.

Suprathreshold detection performance was assessed using reaction times (RT) in Phase 3, in which OAE and EEG were concurrently measured with behavioral detection. As a result of task difficulty equalization, accuracy, and sensitivity measures were no longer informative in highlighting distinctions in the detection of deviants at varying positions. Participants generally showed high accuracy (mean ± 1SEM) for all three positions (early: 96.4 ± 2.0%; middle:95.3 ± 2.0%; late:96.1 ± 2.4%) with no statistically significant difference F [2, 40] = 0.2, p = 0.806). This result further confirms the effectiveness of task equalization using equal SL deviants across positions. To use RT meaningfully, we fitted an exponential Gaussian function to individual RT data (see example in Fig 2H) and extracted three parameters from the fitted distribution (Lacouture and Cousineau, 2008): mean (mu) and standard deviation (sigma) of the Gaussian component, and the exponential component (tau). Individual data and group trends for these parameters are plotted in Figs 2E-2G. The effect of temporal positions on the mean RT was significant (Fig 2E;F [2, 39] = 3.3, p = 0.046). The mean RT for the early position was significantly larger, i.e., slower, than that for the middle position (t (1, 42.1) = 2.5, p = 0.044) but not significantly different from that for the late position (t (1, 42.1) = 1.6, p = 0.263). No significant effect of temporal position on the standard deviation of RT (Fig 2F, F [2, 39] = 0.5, p = 0.592) and the mean of the exponential component (Fig 2G, F [2, 39] = 0.1, p = 0.870) was found. The Gaussian component of the RT is to be interpreted as a reflection of the summed time taken by the sensory process and the motor act of button pressing and the exponential component as a reflection of the decision-making process (Luce, 1986). Therefore, the observed patterns in the RT parameters may indicate a comparable decision-making process in all positions and a somewhat slowed sensory and/or motor process for the early position. It should be emphasized that these statistical differences in behavioral performance are noted despite equalizing task difficulty across the three positions.

### 3.3 Efferent (MOCR) activity

OAE amplitudes were measured concurrently with the behavioral detection of the intensity increment in Phase 3 to characterize the magnitude of MOCR activation. The characteristic exponential decay of the MOCR (group average data plotted in Fig 3A) shows that the efferent activation is significantly different at all three temporal positions, with the least activation at the early position, partial activation at the middle position, and the strongest activation at the late position. The magnitude of reduction in OAE amplitudes (ΔOAE) at each of middle vs. late position, relative to the amplitude at early position, are plotted in Fig 3D (note that the early position was used as a reference for the time course, i.e., time zero). A larger reduction in the OAE amplitude reflects a stronger inhibitory effect of the MOCR (Liberman et al., 1996; Boothalingam et al., 2021). The OAE paradigm used in this study allowed for quantifying the time course of the OAE reduction, making it a useful tool for assessing the relationship between behavioral performance and MOCR activity occurring at a similar temporal position. Raw OAE amplitudes measured at all temporal positions are shown in Fig 3C. The effect of temporal positions on the raw OAE amplitude was significant (F [2, 40] = 79.1, p < 0.001), as expected. The OAE amplitudes measured at the middle and late positions significantly decreased from that measured at the early position (early vs. middle, t (1, 42.1) = 8.8, p < 0.001; early vs. late, t (1, 42.1) = 11.8, p < 0.001; middle vs. late, t (1, 42.1) = 3.0, p= 0.011), indicating effective MOCR activation at the middle and late positions.

**Fig 3.**
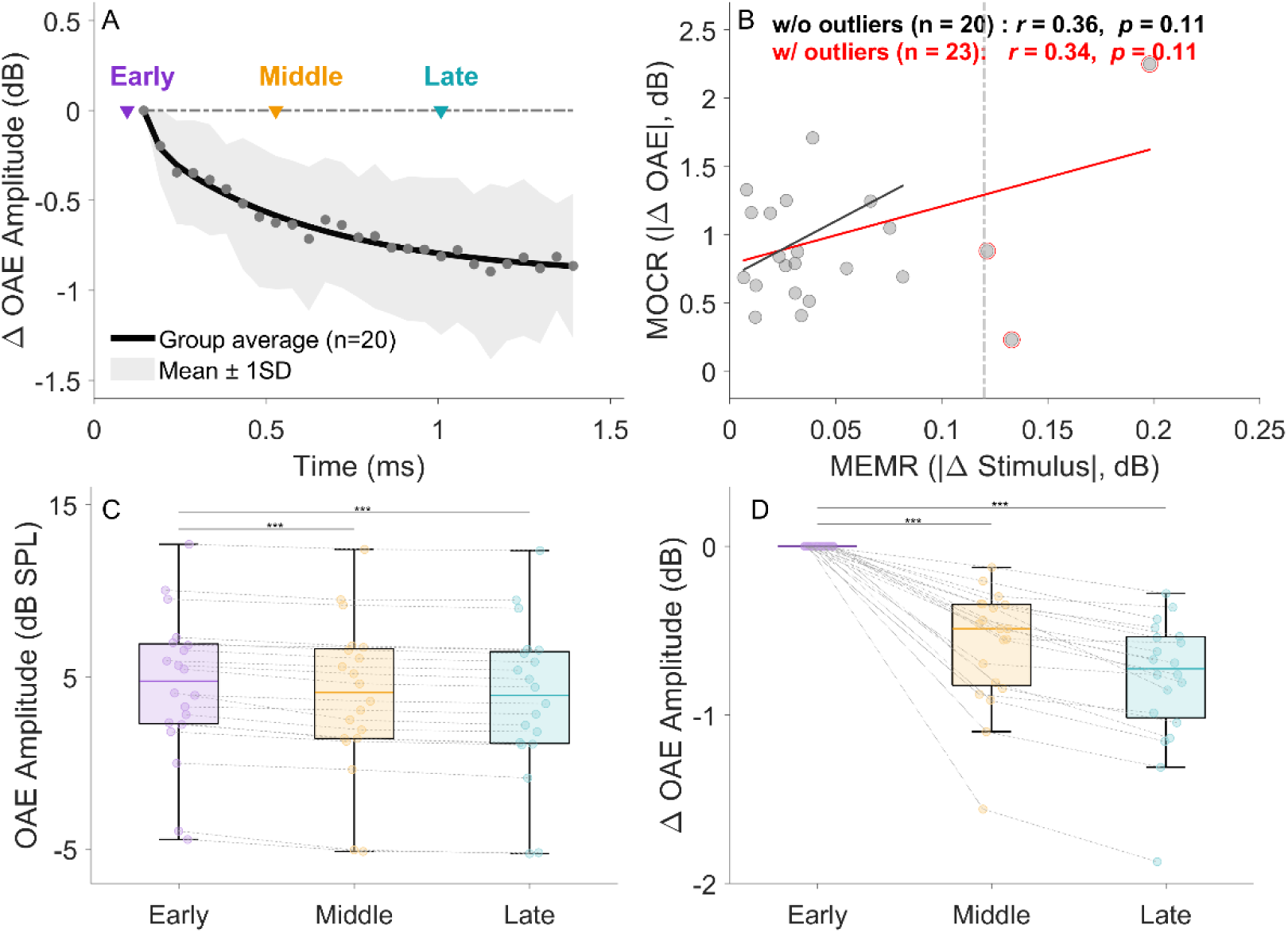
OAE responses evoked by standard click trains. Changes in OAE amplitude from the baseline (time zero) are plotted as the group average in Panel A. The surrounding shaded area represents the group average ± 1 standard deviation. The upside-down triangles mark the temporal position where the deviant triad clicks were imposed in the click train. These temporal positions well captured the null-, partial-, and full-activation of the MOCR, as predicted in Fig 1E. Non-significant correlation between the MOCR and MEMR in 25 participants (*p* = 0.08) is shown in Panel B. Three participants breached the 0.12 dB MEMR threshold (circled in red) and, therefore excluded from all following analyses. The raw OAE data (n = 20) and relative changes in OAEs (re: time zero, i.e., early position) are shown in panels C-D.

### 3.4 Cortical activity

The cortical ACC was reliably elicited by deviants at all three positions (Figs 4A-4C). The he amplitude and latency of the ACC N1-P2 components are plotted in Figs 4D-4F. The effect of temporal position on the summed N1-P2 amplitude was not significant (Fig 4F, F [2, 38] = 1.8, p = 0.179) but the position effect on the N1 latency (Fig 4D, F [2, 37] = 88.1, p < 0.001) and P2 latency (Fig 4E, F [2, 48] = 69.4, p < 0.001) were both significant. Post hoc tests indicated that the significant main effects were driven by the difference between the early and middle positions (N1 latency: t (1, 40.3) = 10.8, p < 0.001; P2 latency: t (1, 41.1) = 9.4, p < 0.001) and between the early and late positions (N1 latency: t (1, 40.3) = 11.4, p < 0.001; P2 latency: t (1, 41.6) = 10.3, p < 0.001). The latency difference between the middle and late positions was not significant (N1 latency: t (1, 40.5) = 0.7, p = 0.783; P2 latency: t (1, 41.6) = 1.1, p = 0.522). N1 latencies (mean ± 1SD) for all three positions were 172.9 ± 22.0 ms, 134.8 ± 18.3 ms, and 134.1 ± 15.0 ms, respectively, which yields a 28.3 to 29.0% faster cortical processing for the middle- and late-positioned deviants relative to the early position. P2 latencies (mean ± 1SD) for all three positions were 258.5 ± 22.3 ms, 213.9 ± 15.5 ms, and 207.8 ± 14.9 ms, respectively, which yields a 20.9 to 24.4% faster cortical processing for the middle- and late-positioned deviants relative to the early position. These results indicate earlier cortical registration of the deviants for the middle and late positions and are in congruence with the improved behavioral detection of deviants (Korczak et al., 2005; Harris et al., 2007).

**Fig 4.**
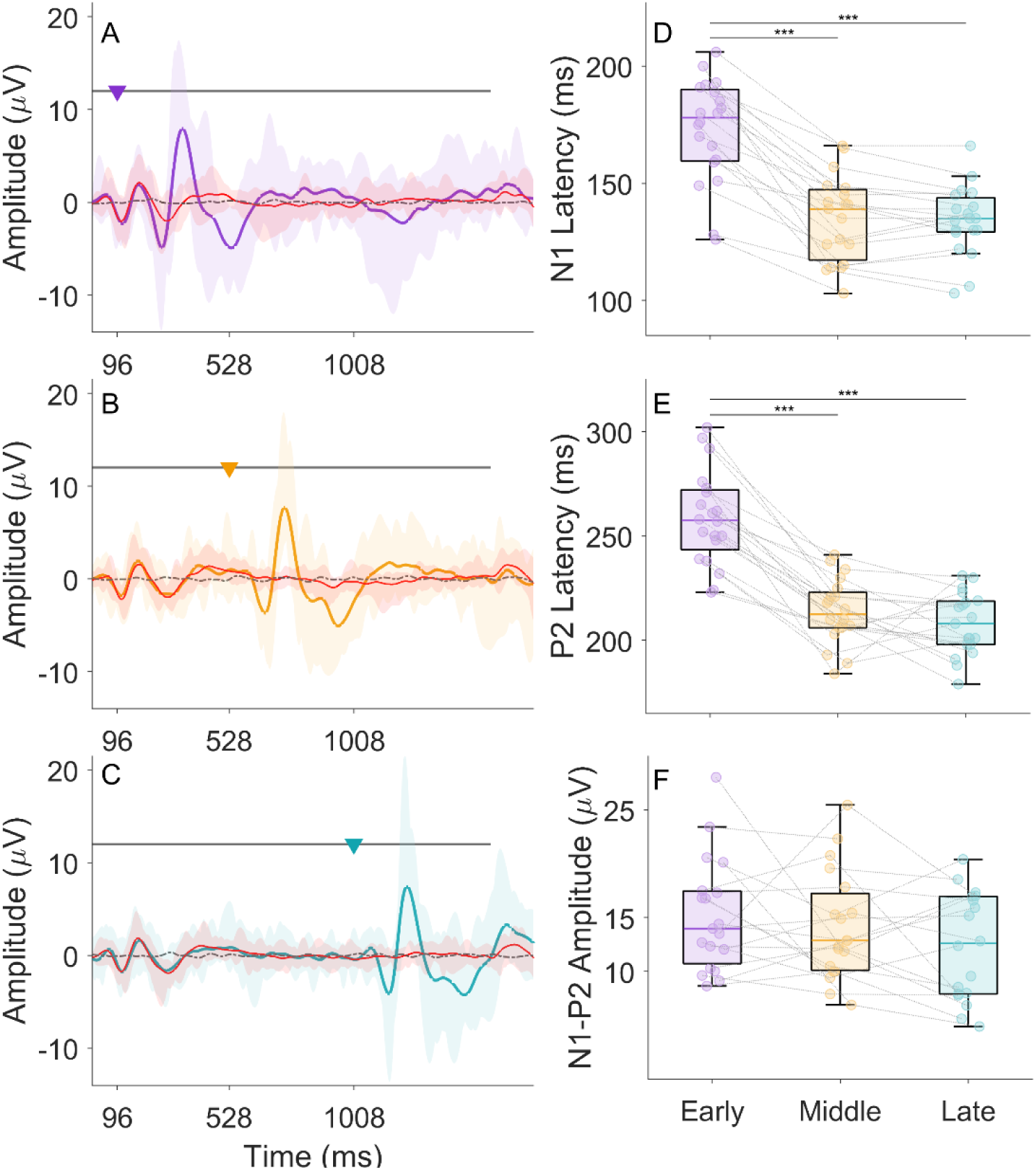
ACC elicited by deviants at three temporal positions. Grand average waveforms across participants (n = 20), plotted for three positions in panels A-C. The upside-down triangles mark the time of onset of deviant click triads in the click trains. EEG response to the standard click trains is plotted in red solid lines in A-C whereas the response to the deviant click trains is plotted in color-coded traces (purple, orange, and cyan for early-, middle-, and late-positioned deviants, respectively). Shaded areas represent the range (minimum to maximum) of individual responses. The dash-dotted line in grey represents the noise floor level in the EEG data. Panels D-F plot the amplitude and latencies for the N1 and P2 components. N1-P2 amplitude was estimated by the peak-to-peak amplitude.

### 3.5 The ear-brain-behavior link: relationship among concurrent measures

The relationships among behavioral, cortical intensity change detection, and MOCR activity were examined using linear mixed-effects regression (LMM) models with repeated measures features. The results of this analysis are shown in Fig 5. Individual data are plotted with fitted lines in different colors. Data for different temporal positions were averaged within each individual and are represented by filled circles on the plot.

**Fig 5.**
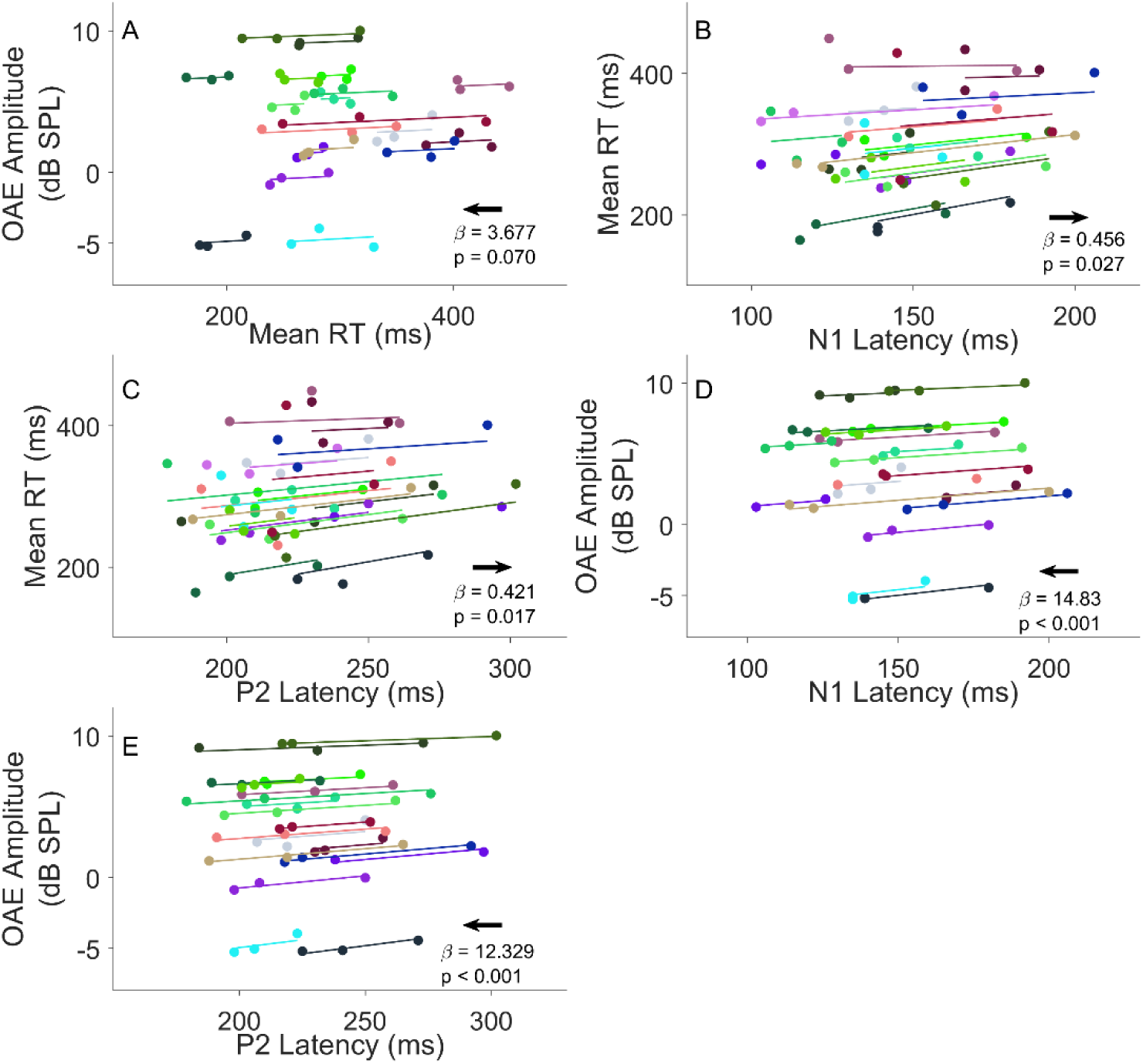
Repeated-measures linear regression results. Repeated-measures LMM models with random intercepts and random slopes were used to assess the relationship among all concurrent measures. Colored, filled circles connected by solid lines are data for all three positions in individual participants (n = 20). All regressions were found statistically significant. Note that the OAE data shown here are the raw OAEs. Therefore, the general pattern of the fitted lines in (A), (D), and (E) indicates that a reduction from the baseline OAE (data with the highest OAE amplitude) is associated with a faster RT or a shorter P2 latency. This trend is indicated by the black right-to-left arrow, which suggests that MOCR activation is related to faster behavioral and cortical detection of the deviants. (B) and (C) reveal a significant positive association between the RT and N1/P2 latency, which indicates that faster behavioral detection is associated with faster cortical detection.

First, the relationship between the raw OAE amplitude, and the mean RT (fixed effect) measured in three temporal positions was investigated. As shown in Figs 3C and 3D, baseline OAE was measured for the early-positioned deviants and had the highest amplitude. The reduction in the OAE amplitude observed for the middle- and late-positioned deviants is an index of MOCR activation strength. The results suggest a positive trend, i.e., a reduction in the OAE amplitude (from the right [early position] to left [late position]) although the association is not statistically significant (Fig 5A, *β* = 3.7, *p* = 0.07, 95% CI [-0.99, 7.586], *η^2^* = 0.08).

Next, the relationship between mean RT and cortical ACC latency (fixed effect) was assessed. Figs 5B and 5C show the statistically significant positive associations between the RT and cortical ACC latency (Fig 5B, *β* = 0.5, *p* = 0.03, 95% CI [0.03, 0.87], *η^2^* = 0.12; Fig 5C, *β* = 0.42, *p* = 0.02, 95% CI [0.08, 0.81], *η^2^* = 0.13). This finding suggests that faster RT is related to earlier cortical registration of deviants.

Finally, the relationship between the raw OAE amplitude and cortical ACC latency (fixed effect) was investigated. Individual data in Figs 5D and 5E show a statistically significant positive correlation suggesting larger MOCR is associated with shorter latency (Fig 5D, N1 latency; *β* = 14.8, *p* < 0.001, 95% CI [10.61, 18.68], *η^2^* = 0.58; Fig 5E, P2 latency; *β* = 12.3, *p* < 0.001, 95% CI [8.49, 16.06], *η^2^* = 0.56). Specifically, in Figs 5D and E, a large, i.e., slow cortical latency is associated with a large OAE raw amplitude (i.e., with the earliest-positioned deviants), and a small, i.e., fast cortical latency is associated with a small OAE raw amplitude (i.e., for the middle and late positions).

Taken together, all concurrent measures with the exception of RT and OAEs are associated with one another, that is, faster behavioral detection is associated with earlier occurring cortical responses and larger MOCR activation. These findings indicate that MOCR activation may play a facilitatory role in expediting cortical registration of a transient acoustic change that may, in turn, lead to a tendency for faster behavioral detection of intensity changes in ongoing sound.

## 4. Discussion

Using a simple yet relevant-to-speech perception task combined with concurrent cochlear, cortical, and behavioral measures, we provide evidence that may suggest the direct association between the MOCR and cortical registration of intensity change detection — an elementary perceptual skill necessary for complex listening, including speech perception in noise.

### 4.1 Potential MOCR association with behavioral intensity change detection

We found lower intensity change detection thresholds for the later-positioned deviants (Fig 2A). The averaged detection threshold for the two later-positioned deviants was about 3.58 dB DSR, which is in line with data previously reported for detecting brief local intensity increments in narrow bands of noises (Grose and Hall, 1997; Moore and Peters, 1997). Grose and Hall (1997) showed that the just-noticeable intensity increment was 3.8 dB when detecting brief local increments in narrow bands of noise. Moore and Peters (1997) reported a higher detection threshold for the intensity increment occurring earlier in a sinusoid, as opposed to later, which was then interpreted as a possible consequence of neural adaptation that is subtractive or inhibitory. The MOCR activity could be one mechanism for such subtractive neural adaptation, leading to a reduction in neural response to the ongoing signal over time while increasing the contrast between the level changes and the ongoing carrier sound. While other adaptive mechanisms such as mean level adaptation (Dean et al., 2005) and contrast adaptation (Rabinowitz et al., 2013) can also play a role in improved detection of later occurring deviants, the extent to which the MOCR contributes to these adaptive mechanisms is not fully understood. Computational auditory periphery modeling of efferent effects at the periphery suggests both the MOCR and the MEMR activity contribute to adaptive processes observed elsewhere in the auditory system (Grange et al., 2022), especially considering mean level adaptation has been reported as early in the auditory system as the auditory nerve (Wen et al., 2012; Willmore and King, 2023). As such, either via adaptive mechanisms or due to a direct anti-masking effect at the periphery (Lauer et al., 2021; Guinan, 2014, 2018; Lopez-Poveda, 2018; Winslow and Sachs, 1987, 1988), the MOCR contributions to improved detection at later positions seem likely.

To our knowledge, very few previous behavioral studies evaluating a MOCR role in perception have used RT measures (Smith and Cone, 2015; Mertes et al., 2019). Because RT measures can be largely confounded by individual differences in task difficulty and performance (McGarrigle et al., 2014; Alhanbali et al., 2019), task difficulty and performance must be equalized, as we did in the present study (Houben et al., 2013; Strand et al., 2018; Alhanbali et al., 2019). The facilitative effect of the MOCR is evident in faster behavioral intensity change detection for later occurring deviants (Fig 2E), despite equalizing task difficulty and detection accuracy across deviant positions (Figs. 2C and 2D). As such, the improvement in behavioral detection is unlikely to stem from any acoustic features, including energy or salience differences between deviant positioning within the stimulus, and is more likely to be related to underlying physiological responses, at least partly influenced by the MOCR. As such, the stronger MOCR activity at the later temporal positions may have played a role in the ease of perception and decision-making processes at these positions.

Previous investigations attempting to link overshoot-like effects in OAE measures and the psychophysical effects in the same listeners have produced mixed results (Keefe et al., 2009; Walsh et al., 2010; Henin and Long, 2013). These studies measured the change in the OAE responses to a tonal signal embedded in the early or late portions of a noise masker. A similar overshoot pattern in both OAEs and detection thresholds would be considered the result of an underlying association between the MOCR mediation of the cochlear gain and the behavioral detection (Walsh et al., 2010). However, such correlational analysis are not currently available in the literature. The click train paradigm used in this study differs from the classical overshoot paradigm in that the deviant triad signal shares the same acoustic characteristics (except the level) with the ongoing background click train — both were clicks. The contrast between the deviant triad and the standard click train ongoing in the background is relatively smaller than that between a tone and a noise. This makes the detection task in this study more challenging. Here, we present evidence suggesting the role of the MOCR likely facilitating a trend of earlier behavioral detection of the deviant, but we also acknowledge other neural phenomena, both in the afferent and efferent pathways, may play further roles.

### 4.2 MOCR activation is associated with cortical intensity change detection

We found that the N1-P2 ACC complex was evoked significantly earlier (by 20-29%) by intensity changes in the later positions relative to the early position. Together with the significant correlation between ACC latency and MOCR activation, our results indicate that the MOCR likely facilitates earlier cortical registration of intensity changes. This is in line with previous ACC studies that, using intensity discrimination tasks, demonstrated improved behavioral discriminability of intensity change associated with shorter ACC latency (Dimitrijevic et al., 2009; He et al., 2012). In addition to serving as a secondary validation to MOCR facilitation of behavioral intensity change detection, the MOCR-ACC latency relationship suggests that at least the pre-attentive obligatory responses in the cortex may be influenced by MOCR activity and potentially other feedback networks (Hernández-Pérez et al., 2021).

Although the ACC amplitude increases with greater acoustic changes (Martin and Boothroyd, 2000; Harris et al., 2007; Dimitrijevic et al., 2009, 2011; He et al., 2012), the lack of significant difference in ACC amplitudes across the three positions is not unexpected as the deviants (intensity changes) were presented at suprathreshold equal sensation level. This lack of significant amplitude difference between positions adds further authentication to MOCR facilitation of behavioral and cortical responses as there are no discernable acoustical differences between deviants at these positions, which primarily drive the amplitude of the N1-P2 ACC (Martin and Boothroyd, 2000; Harris et al., 2007; Dimitrijevic et al., 2009; He et al., 2012). That is, the MOCR facilitation in processing speed occurs despite the lack of any acoustically perceptible, and cortical activation amplitude, differences.

How may the MOCR reduction in cochlear amplification contribute to faster cortical registration of deviants? There are no definitive answers for finding; animal work that investigates the MOCR concurrently with cortical and subcortical neural activity may shed light on a potentially causal relationship. However, two speculations could be made. (1) The improvement in transient detection could also be thought of as a cleaner, less jittered, input to the cochlear nucleus. That is, an improvement in SNR as described by the MOCR unmasking hypothesis (Guinan, 2006; Winslow and Sachs, 1987). While it is well known that the cochlear nucleus cells refine ‘jittery’ auditory nerve spikes (Joris et al., 1994), more salient, i.e., less jitter in spiking of the auditory nerve could be expected to further increase the probability of spiking in the cochlear nucleus neurons. This increase in the spiking confidence, potentially arising from an increase in coincident spiking, could be thought to carry over to later stages and contribute to a more robust signal leading to earlier spiking in higher-level auditory centers. This effect of the MOCR could be thought of a leftward shift in the probability density function of neuronal spiking in the cochlear nucleus and beyond. (2) The MOCR may advance the phase of the cochlear output, which may be carried over, and possibly further refined in later stages of the auditory system, especially pertaining to binaural fusion. There is evidence for the lateral OC (Darrow et al., 2006) and a suggestion for an MEMR (Cho et al., 2023) role in binaural processing, and therefore an MOCR role is also conceivable.

### 4.3 Behavioral intensity change detection is associated with cortical activation latency

The ACC is sensitive to intensity changes as small as 2 dB and is therefore considered an effective metric for detection threshold estimation. Detection and discrimination thresholds estimated by ACC amplitude are comparable with thresholds measured psychophysically in young (Martin and Boothroyd, 2000; Dimitrijevic et al., 2009; He et al., 2012) and older listeners (Harris et al., 2007). The results from the present study are most consistent with this line of research but are different in that a significant correlation was found between behavioral performance and the ACC latency (Fig 4D) as opposed to ACC amplitude. This is largely due to the design of the paradigm, where we equalized the detectability of intensity changes across the three temporal positions. As a result, the detection accuracy-based behavioral measures were statistically indistinguishable. Therefore, the main comparison was made between RT and the ACC measures. The association between smaller ACC latencies (Figs 4B and 4C) were associated with smaller RTs, suggest that earlier cortical registration of the deviant, i.e., the intensity change, likely subserves faster behavioral detection.

### 4.4 No influence of the middle ear muscle reflex

If the MEMR is activated alongside the MOCR, it might also be involved in the down-regulation of signals admitted into the system, albeit by altering the middle ear impedance characteristics. This is a possible confounding factor in all MOCR studies (Guinan, 2014b; Feeney et al., 2017; Xu et al., 2017; Lopez-Poveda, 2018). Based on the analysis of click stimuli recorded in the ear canal, we rejected 3 participants who demonstrated >0.12 dB change in the stimulus - a potential indicator of MEMR activation. If the MEMR activation was driving the changes in the OAEs reported herein, we would expect to see a positive correlation between stimulus and OAE changes over time (1s). We did not see any indication of such a correlation (Fig 3B). These quality checks notwithstanding, we performed further analyses to assess the possible effects of MEMR on improved behavioral and cortical detections. Unlike the relationships observed with OAE changes (MOCR), no significant associations between the MEMR strength and the RTs (*β* = −0.01, *p* = 0.95, 95% CI [-0.22, 0.21], *η^2^* = 0), or between the MEMR strength and the ACC latencies (N1 latency: *β* = 0.27, *p* = 0.95, 95% CI [-0.22, 0.21], *η^2^* = 0; P2 latency: *β* = 0.18, *p* = 0.16, 95% CI [-0.06, 0.41], *η^2^* = 0.05) were observed. These results support that the deviant position effect in the behavioral and cortical responses reported in this study is driven by the MOCR and not the MEMR.

### 4.5 Implications for speech perception

Elementary acoustic features that eventually combine to generate a speech percept are dynamic and distributed across cortical and subcortical areas in a complex feedforward and feedback network (Rauschecker and Scott, 2009; Alluri et al., 2012; Bidelman et al., 2019). More importantly, these features appear invariant to subtle changes in the acoustics (Formisano et al., 2008; Leaver and Rauschecker, 2010). Such robustness is critical for protection against corruption from noise and signal distortion. Further, we now also know that multiple feedback loops in the subcortical regions and the cortex facilitate auditory perception differently (Hernández-Pérez et al., 2021).

Some of the robustness noted in speech might be a result of feedback systems, such as the MOCR, dynamically improving SNR in the system (Oertel et al., 2011; Parbery-Clark et al., 2011; Farhadi et al., 2021; Boothalingam et al., 2023; Hockley et al., 2022). We recently showed that the MOCR inhibition of cochlear activity in humans is compensated in the brainstem through local feedback loops (Boothalingam et al., 2023). Animal work have posited this feedback loop likely facilitates complex signal (e.g., speech) encoding by boosting spectral peaks and thereby improving the signal-to-noise ratio (Fujino and Oertel, 2001; Mulders et al., 2002, 2007; Hockley et al., 2022). Mathematical models suggest MOCR activation improves firing in the inferior colliculus for amplitude-modulated noise, often a proxy for speech stimuli (Farhadi et al., 2021). Such MOCR activity is likely only partially reflected when indexed using OAEs. Similarly, changes inflicted by the MOCR at the periphery and in the brainstem do not necessitate similar and linearly associated changes in cortical areas (Boothalingam et al., 2023). That is, even if the MOCR facilitates the perception of elementary acoustic features in speech, ultimate perception and comprehension of speech may not directly reflect MOCR activity.

Given such complex orchestration of processes across a large and distributed brain network, it is not surprising that previous studies that attempted to correlate speech perception with MOCR activation, especially those obtained separately and independently of each other, report mixed results. To circumvent this caveat, we employed a simpler intensity change detection task and found that the MOCR likley facilitates auditory perception, consistent with previous psychophysical evidence. These findings are relevant because encoding rapid temporal fluctuations is critical to speech perception in noise (Shannon et al., 1995; Turner et al., 1995; Xu and Zheng, 2007; Swaminathan and Heinz, 2012). As such, our results provide evidence for an active role of the MOCR in elementary perception that likely contributes to speech perception despite the mixed findings from past human studies. Understanding the role of auditory efferents in speech perception may help devise better therapeutics (Le Prell et al., 2003; Liberman et al., 2014; Boero et al., 2018, 2020) and technology (Liu and Demosthenous, 2020; Farhadi et al., 2021) for treating hearing damage.

Overall, our findings corroborate previous human psychophysical data that suggests an active role of the MOCR in human signal detection and discrimination. Specifically, our data shows facilitated detection in the later parts of a click train, where the MOCR activation is stronger, indicated by faster detection and greater sensitivity, and earlier cortical registration when task difficulty and performance are equalized. This demonstration of novel findings is owed to our paradigm where we measured cochlear, cortical, and behavioral responses concurrently.

## Acknowledgments

The authors thank Veronika Mak for her help with data acquisition. This study was supported by funds from the Office of the Vice Chancellor for Research and Graduate Education, University of Wisconsin-Madison, to S.B.

## References

1. Abdala C, Mishra S, Garinis A (2013) Maturation of the human medial efferent reflex revisited. The Journal of the Acoustical Society of America 133:938–950.

2. Alhanbali S, Dawes P, Millman RE, Munro KJ (2019) Measures of Listening Effort Are Multidimensional: Ear and Hearing 40:1084–1097.

3. Alluri V, Toiviainen P, Jääskeläinen IP, Glerean E, Sams M, Brattico E (2012) Large-scale brain networks emerge from dynamic processing of musical timbre, key and rhythm. NeuroImage 59:3677–3689.

4. Backus BC, Guinan JJ (2006) Time-course of the human medial olivocochlear reflex. The Journal of the Acoustical Society of America 119:2889–2904.

5. Bacon SP, Healy EW (2000) Effects of ipsilateral and contralateral precursors on the temporal effect in simultaneous masking with pure tones. The Journal of the Acoustical Society of America 107:1589–1597.

6. Bacon SP, Moore BCJ (1986) Temporal effects in simultaneous pure-tone masking: effects of signal frequency, masker/signal frequency ratio, and masker level. Hearing Research 23:257–266.

7. Bates D, Mächler M, Bolker B, Walker S (2015) Fitting Linear Mixed-Effects Models Using **lme4**. J Stat Soft 67 Available at: http://www.jstatsoft.org/v67/i01/ [Accessed December 19, 2021].

8. Beim JA, Elliott M, Oxenham AJ, Wojtczak M (2015) Stimulus Frequency Otoacoustic Emissions Provide No Evidence for the Role of Efferents in the Enhancement Effect. JARO 16:613–629.

9. Bidelman GM, Price CN, Shen D, Arnott SR, Alain C (2019) Afferent-efferent connectivity between auditory brainstem and cortex accounts for poorer speech-in-noise comprehension in older adults. Hearing Research 382:107795.

10. Boero LE, Castagna VC, Guilmi MND, Goutman JD, Elgoyhen AB, Gómez-Casati ME (2018) Enhancement of the Medial Olivocochlear System Prevents Hidden Hearing Loss. J Neurosci 38:7440–7451.

11. Boero LE, Castagna VC, Terreros G, Moglie MJ, Silva S, Maass JC, Fuchs PA, Delano PH, Elgoyhen AB, Gómez-Casati ME (2020) Preventing presbycusis in mice with enhanced medial olivocochlear feedback. Proc Natl Acad Sci USA 117:11811–11819.

12. Boothalingam S, Easwar V, Bross A (2022) External and middle ear influence on envelope following responses. The Journal of the Acoustical Society of America 152:2794–2803.

13. Boothalingam S, Goodman SS (2021) Click evoked middle ear muscle reflex: Spectral and temporal aspects. The Journal of the Acoustical Society of America 149:2628–2643.

14. Boothalingam S, Goodman SS, MacCrae H, Dhar S (2021) A Time-Course-Based Estimation of the Human Medial Olivocochlear Reflex Function Using Clicks. Frontiers in Neuroscience 15 Available at: https://www.frontiersin.org/articles/10.3389/fnins.2021.746821 [Accessed February 5, 2023].

15. Boothalingam S, Peterson A, Powell L, Easwar V (2023) Auditory brainstem mechanisms likely compensate for self-imposed peripheral inhibition. Sci Rep 13:12693.

16. Capps MJ, Ades HW (1968) Auditory frequency discrimination after transection of the olivocochlear bundle in squirrel monkeys. Experimental Neurology 21:147–158.

17. Castro-Palacio JC, Fernández-de-Córdoba P, Isidro JM, Sahu S, Navarro-Pardo E (2021) Human Reaction Times: Linking Individual and Collective Behaviour Through Physics Modeling. Symmetry 13:451.

18. Cho NH, Ravicz ME, & Puria S (2023) Human middle-ear muscle pulls change tympanic-membrane shape and low-frequency middle-ear transmission magnitudes and delays. Hearing Research 430: 108721.

19. Clause A, Lauer AM, Kandler K (2017) Mice Lacking the Alpha9 Subunit of the Nicotinic Acetylcholine Receptor Exhibit Deficits in Frequency Difference Limens and Sound Localization. Frontiers in Cellular Neuroscience 11 Available at: https://www.frontiersin.org/articles/10.3389/fncel.2017.00167 [Accessed September 5, 2023].

20. Darrow KN, Maison SF, & Liberman MC (2006) Cochlear efferent feedback balances interaural sensitivity. Nature neuroscience 9(12): 1474–1476.

21. de Boer E (1986) On Thresholds of Short-Duration Intensity Increments and Decrements. In: Auditory Frequency Selectivity (Moore BCJ, Patterson RD, eds), pp 429–436. Boston, MA: Springer US. Available at: http://link.springer.com/10.1007/978-1-4613-2247-4_46 [Accessed August 4, 2022].

22. Dean I, Harper NS, McAlpine D (2005) Neural population coding of sound level adapts to stimulus statistics. Nat Neurosci 8:1684–1689.

23. Delano PH, Elgueda D, Hamame CM, Robles L (2007) Selective Attention to Visual Stimuli Reduces Cochlear Sensitivity in Chinchillas. J Neurosci 27:4146–4153.

24. Dewson JH (1968) Efferent Olivocochlear bundle: some relationships to stimulus discrimination in noise. Journal of Neurophysiology 31:122–130.

25. Dimitrijevic A, Lolli B, Michalewski HJ, Pratt H, Zeng F-G, Starr A (2009) Intensity changes in a continuous tone: Auditory cortical potentials comparison with frequency changes. Clinical Neurophysiology 120:374–383.

26. Dimitrijevic A, Starr A, Bhatt S, Michalewski HJ, Zeng F-G, Pratt H (2011) Auditory cortical N100 in pre- and post-synaptic auditory neuropathy to frequency or intensity changes of continuous tones. Clinical Neurophysiology 122:594–604.

27. Dolan DF, Nuttall AL (1988) Masked cochlear whole-nerve response intensity functions altered by electrical stimulation of the crossed olivocochlear bundle. The Journal of the Acoustical Society of America 83:1081–1086.

28. Elliott LL (1965) Changes in the Simultaneous Masked Threshold of Brief Tones. The Journal of the Acoustical Society of America 38:738–746.

29. Farhadi A, Jennings SG, Strickland EA, Carney LH (2021) A Closed-Loop Gain-Control Feedback Model for The Medial Efferent System of The Descending Auditory Pathway. In: ICASSP 2021 - 2021 IEEE International Conference on Acoustics, Speech and Signal Processing (ICASSP), pp 291–295.

30. Feeney MP, Keefe DH, Hunter LL, Fitzpatrick DF, Garinis AC, Putterman DB, McMillan GP (2017) Normative Wideband Reflectance, Equivalent Admittance at the Tympanic Membrane, and Acoustic Stapedius Reflex Threshold in Adults: Ear and Hearing 38:e142–e160.

31. Fletcher MD, Krumbholz K, de Boer J (2016) Effect of Contralateral Medial Olivocochlear Feedback on Perceptual Estimates of Cochlear Gain and Compression. JARO 17:559–575.

32. Formisano E, De Martino F, Bonte M, Goebel R (2008) “Who” Is Saying “What”? Brain-Based Decoding of Human Voice and Speech. Science 322:970–973.

33. Fujino K, Oertel D (2001) Cholinergic Modulation of Stellate Cells in the Mammalian Ventral Cochlear Nucleus. J Neurosci 21:7372–7383.

34. Geisler CD (1974) Model of crossed olivocochlear bundle effects. The Journal of the Acoustical Society of America 56:1910–1912.

35. Grange J, Zhang M, Culling J (2022) The Role of Efferent Reflexes in the Efficient Encoding of Speech by the Auditory Nerve. J Neurosci 42:6907–6916.

36. Grose JH, Hall JWH (1997) Multiband detection of energy fluctuations. The Journal of the Acoustical Society of America 102:1088–1096.

37. Guinan JJ (2006) Olivocochlear Efferents: Anatomy, Physiology, Function, and the Measurement of Efferent Effects in Humans: Ear and Hearing 27:589–607.

38. Guinan JJ (2014a) Cochlear Mechanics, Otoacoustic Emissions, and Medial Olivocochlear Efferents: Twenty Years of Advances and Controversies Along with Areas Ripe for New Work. In: Perspectives on Auditory Research (Popper AN, Fay RR, eds), pp 229–246 Springer Handbook of Auditory Research. New York, NY: Springer. Available at: 10.1007/978-1-4614-9102-6_13 [Accessed February 14, 2023].

39. Guinan JJ (2014b) Olivocochlear efferent function: issues regarding methods and the interpretation of results. Front Syst Neurosci 8 Available at: http://journal.frontiersin.org/article/10.3389/fnsys.2014.00142/abstract [Accessed March 26, 2020].

40. Guinan JJ (2018) Olivocochlear efferents: Their action, effects, measurement and uses, and the impact of the new conception of cochlear mechanical responses. Hearing Research 362:38–47.

41. Harris KC, Mills JH, Dubno JR (2007) Electrophysiologic correlates of intensity discrimination in cortical evoked potentials of younger and older adults. Hearing Research 228:58–68.

42. Hautus MJ (1995) Corrections for extreme proportions and their biasing effects on estimated values ofd′. Behavior Research Methods, Instruments, & Computers 27:46–51.

43. He S, Grose JH, Buchman CA (2012) Auditory discrimination: The relationship between psychophysical and electrophysiological measures. International Journal of Audiology 51:771–782.

44. Heinz MG, Formby C (1999) Detection of time- and bandlimited increments and decrements in a random-level noise. The Journal of the Acoustical Society of America 106:313–326.

45. Henin S, Long GR (2013) Evaluating the role of efferent inhibition on cochlear responses: Simultaneous psychophysical and otoacoustic emission measurements. Proceedings of Meetings on Acoustics 19:050092.

46. Hernández-Pérez H, Mikiel-Hunter J, McAlpine D, Dhar S, Boothalingam S, Monaghan JJM, McMahon CM (2021) Understanding degraded speech leads to perceptual gating of a brainstem reflex in human listeners. PLOS Biology 19:e3001439.

47. Hicks ML, Bacon SP (1992) Factors influencing temporal effects with notched-noise maskers. Hearing Research 64:123–132.

48. Hienz RD, Stiles P, May BJ (1998) Effects of bilateral olivocochlear lesions on vowel formant discrimination in cats. Hearing Research 116:10–20.

49. Hockley A, Wu C, Shore SE (2022) Olivocochlear projections contribute to superior intensity coding in cochlear nucleus small cells. The Journal of physiology 600(1): 61–73.

50. Houben R, van Doorn-Bierman M, Dreschler WA (2013) Using response time to speech as a measure for listening effort. International Journal of Audiology 52:753–761.

51. James AL, Harrison RV, Pienkowski M, Dajani HR, & Mount RJ (2005) Dynamics of real time DPOAE contralateral suppression in chinchillas and humans. International journal of audiology 44(2):118–129.

52. Jennings SG, Heinz MG, Strickland EA (2011) Evaluating Adaptation and Olivocochlear Efferent Feedback as Potential Explanations of Psychophysical Overshoot. J Assoc Res Otolaryngol 12:345–360.

53. Jennings SG, Strickland EA (2012) Evaluating the effects of olivocochlear feedback on psychophysical measures of frequency selectivity. The Journal of the Acoustical Society of America 132:2483–2496.

54. Joris PX, Carney LH, Smith PH, & Yin TC (1994) Enhancement of neural synchronization in the anteroventral cochlear nucleus. I. Responses to tones at the characteristic frequency. Journal of neurophysiology 71(3): 1022–1036.

55. Kawase T, Delgutte B, Liberman MC (1993) Antimasking effects of the olivocochlear reflex. II. Enhancement of auditory-nerve response to masked tones. Journal of Neurophysiology 70:2533–2549.

56. Kawase T, Liberman MC (1993) Antimasking effects of the olivocochlear reflex. I. Enhancement of compound action potentials to masked tones. Journal of Neurophysiology 70:2519–2532.

57. Keefe DH, Schairer KS, Ellison JC, Fitzpatrick DF, Jesteadt W (2009) Use of stimulus-frequency otoacoustic emissions to investigate efferent and cochlear contributions to temporal overshoot. The Journal of the Acoustical Society of America 125:1595–1604.

58. Kim DO, Dorn PA, Neely ST, Gorga MP (2001) Adaptation of Distortion Product Otoacoustic Emission in Humans. J Assoc Res Otolaryngol 2:31–40.

59. Korczak PA, Kurtzberg D, Stapells DR (2005) Effects of Sensorineural Hearing Loss and Personal Hearing Aids on Cortical Event-Related Potential and Behavioral Measures of Speech-Sound Processing. Ear and Hearing 26:165.

60. Kumar UA, Vanaja CS (2004) Functioning of Olivocochlear Bundle and Speech Perception in Noise: Ear and Hearing 25:142–146.

61. Kuznetsova A, Brockhoff PB, Christensen RHB (2017) lmerTest Package: Tests in Linear Mixed Effects Models. Journal of Statistical Software 82:1–26.

62. Lacouture Y, Cousineau D (2008) How to use MATLAB to fit the ex-Gaussian and other probability functions to a distribution of response times. TQMP 4:35–45.

63. Laird NM, Ware JH (1982) Random-Effects Models for Longitudinal Data. Biometrics 38:963–974.

64. Lauer AM, Jimenez SV, Delano PH (2021) Olivocochlear efferent effects on perception and behavior. Hearing Research:108207.

65. Le Prell CG, Shore SE, Hughes LF, Bledsoe SC (2003) Disruption of Lateral Efferent Pathways: Functional Changes in Auditory Evoked Responses. JARO 4:276–290.

66. Leaver AM, Rauschecker JP (2010) Cortical Representation of Natural Complex Sounds: Effects of Acoustic Features and Auditory Object Category. J Neurosci 30:7604–7612.

67. Levine TR, Hullett CR (2002) Eta Squared, Partial Eta Squared, and Misreporting of Effect Size in Communication Research. Human Communication Research 28:612–625.

68. Liberman MC, Guinan JJ (1998) Feedback control of the auditory periphery: anti-masking effects of middle ear muscles vs. olivocochlear efferents. Journal of communication disorders 31:471–483.

69. Liberman MC, Liberman LD, Maison SF (2014) Efferent Feedback Slows Cochlear Aging. J Neurosci 34:4599–4607.

70. Liberman MC, Puria S, Guinan jJ (1996) The ipsilaterally evoked olivocochlear reflex causes rapid adaptation of the 2 f1−f2 distortion product otoacoustic emission. The Journal of the Acoustical Society of America 99:3572–3584.

71. Lilaonitkul W, Guinan JJ (2012) Frequency tuning of medial-olivocochlear-efferent acoustic reflexes in humans as functions of probe frequency. Journal of Neurophysiology 107:1598–1611.

72. Liu F, Demosthenous A (2020) Effect of Time Constant on Speech Enhancement in Hearing Aids Based on Auditory Neural Feedback. In: 2020 IEEE International Symposium on Circuits and Systems (ISCAS), pp 1–5 Available at: https://ieeexplore.ieee.org/abstract/document/9181228 [Accessed January 5, 2024].

73. Lopez-Poveda EA (2018) Olivocochlear Efferents in Animals and Humans: From Anatomy to Clinical Relevance. Front Neurol 9 Available at: https://www.frontiersin.org/articles/10.3389/fneur.2018.00197/full [Accessed March 26, 2020].

74. Lopez-Poveda EA, Aguilar E, Johannesen PT, Eustaquio-Martín A (2013) Contralateral Efferent Regulation of Human Cochlear Tuning: Behavioural Observations and Computer Model Simulations. In: Basic Aspects of Hearing (Moore BCJ, Patterson RD, Winter IM, Carlyon RP, Gockel HE, eds), pp 47–54 Advances in Experimental Medicine and Biology. New York, NY: Springer.

75. Luce RD (1986) Response Times: Their Role in Inferring Elementary Mental Organization. Oxford University Pres, U.S.A.

76. Macmillan NA, Creelman CD (2004) Detection Theory: A User’s Guide. Psychology Press.

77. Marrufo-Pérez MI, Eustaquio-Martín A, López-Bascuas LE, Lopez-Poveda EA (2018) Temporal Effects on Monaural Amplitude-Modulation Sensitivity in Ipsilateral, Contralateral and Bilateral Noise. Journal of the Association for Research in Otolaryngology 19:147–161.

78. Martin BA, Boothroyd A (2000) Cortical, auditory, evoked potentials in response to changes of spectrum and amplitude. The Journal of the Acoustical Society of America 107:2155–2161.

79. Maruthy S, Kumar UA, Gnanateja GN (2017) Functional Interplay Between the Putative Measures of Rostral and Caudal Efferent Regulation of Speech Perception in Noise. JARO 18:635–648.

80. Matzke D, Wagenmakers E-J (2009) Psychological interpretation of the ex-Gaussian and shifted Wald parameters: A diffusion model analysis. Psychonomic Bulletin & Review 16:798–817.

81. McFadden D, Champlin CA (1990) Reductions in overshoot during aspirin use. The Journal of the Acoustical Society of America 87:2634–2642.

82. McGarrigle R, Munro KJ, Dawes P, Stewart AJ, Moore DR, Barry JG, Amitay S (2014) Listening effort and fatigue: What exactly are we measuring? A British Society of Audiology Cognition in Hearing Special Interest Group ‘white paper.’ International Journal of Audiology 53:433–445.

83. Mertes IB, Johnson KM (2020) Lack of association between contralateral inhibition of otoacoustic emissions and vowel formant discrimination in noise. Hearing, Balance and Communication:1–6.

84. Mertes IB, Johnson KM, Dinger ZA (2019) Olivocochlear efferent contributions to speech-in-noise recognition across signal-to-noise ratios. The Journal of the Acoustical Society of America 145:1529–1540.

85. Mertes IB, Wilbanks EC, Leek MR (2018) Olivocochlear Efferent Activity Is Associated With the Slope of the Psychometric Function of Speech Recognition in Noise: Ear and Hearing 39:583–593.

86. Micheyl C, Collet L (1996) Involvement of the olivocochlear bundle in the detection of tones in noise. The Journal of the Acoustical Society of America 99:1604–1610.

87. Micheyl C, Morlet T, Giraud AL, Collet L, Morgon A (1995) Contralateral Suppression of Evoked Otoacoustic Emissions and Detection of a Multi-Tone Complex in Noise. Acta Oto-Laryngologica 115:178–182.

88. Micheyl C, Perrot X, Collet L (1997) Relationship Between Auditory Intensity Discrimination in Noise and Olivocochlear Efferent System Activity in Humans. Behavioral neuroscience 111:801.

89. Moore BCJ, Peters RW (1997) Detection of increments and decrements in sinusoids as a function of frequency, increment, and decrement duration and pedestal duration. The Journal of the Acoustical Society of America 102:2954–2965.

90. Morand-Villeneuve N, Garnier S, Grimault N, Veuillet E, Collet L, Micheyl C (2002) Medial olivocochlear bundle activation and perceived auditory intensity in humans. Physiology & Behavior 77:311–320.

91. Mulders WHAM, Harvey AR, Robertson D (2007) Electrically Evoked Responses in Onset Chopper Neurons in Guinea Pig Cochlear Nucleus. Journal of Neurophysiology 97:3288–3297.

92. Mulders WHAM, Winter IM, Robertson D (2002) Dual action of olivocochlear collaterals in the guinea pig cochlear nucleus. Hearing Research 174:264–280.

93. Oertel D, Wright S, Cao X-J, Ferragamo M, Bal R (2011) The multiple functions of T stellate/multipolar/chopper cells in the ventral cochlear nucleus. Hearing Research 276:61–69.

94. Onishi S, Davis H (1968) Effects of Duration and Rise Time of Tone Bursts on Evoked V Potentials. The Journal of the Acoustical Society of America 44:582–591.

95. Oxenham AJ (1997) Increment and decrement detection in sinusoids as a measure of temporal resolution. The Journal of the Acoustical Society of America 102:1779–1790.

96. Parbery-Clark A, Marmel F, Bair J, Kraus N (2011) What subcortical-cortical relationships tell us about processing speech in noise: Neural correlates of speech in noise. European Journal of Neuroscience 33:549–557.

97. Rabinowitz NC, Willmore BDB, King AJ, Schnupp JWH (2013) Constructing Noise-Invariant Representations of Sound in the Auditory Pathway Zador AM, ed. PLoS Biol 11:e1001710.

98. Rao A, Koerner TK, Madsen B, Zhang Y (2020) Investigating Influences of Medial Olivocochlear Efferent System on Central Auditory Processing and Listening in Noise: A Behavioral and Event-Related Potential Study. Brain Sciences 10:428.

99. Rasetshwane DM, Neely ST (2011) Calibration of otoacoustic emission probe microphones. The Journal of the Acoustical Society of America 130:EL238–EL243.

100. Rauschecker JP, Scott SK (2009) Maps and streams in the auditory cortex: nonhuman primates illuminate human speech processing. Nat Neurosci 12:718–724.

101. Rodríguez J, Neely ST, Jesteadt W, Tan H, Gorga MP (2011) Comparison of distortion-product otoacoustic emission growth rates and slopes of forward-masked psychometric functionsa). The Journal of the Acoustical Society of America 129:864–875.

102. Roverud E, Strickland EA (2010) The time course of cochlear gain reduction measured using a more efficient psychophysical techniquea). The Journal of the Acoustical Society of America 128:1203–1214.

103. Roverud E, Strickland EA (2014) Accounting for nonmonotonic precursor duration effects with gain reduction in the temporal window modela). The Journal of the Acoustical Society of America 135:1321–1334.

104. Schairer KS, Messersmith J, Jesteadt W (2008) Use of psychometric-function slopes for forward-masked tones to investigate cochlear nonlinearitya). The Journal of the Acoustical Society of America 124:2196–2215.

105. Schairer KS, Nizami L, Reimer JF, Jesteadt W (2003) Effects of peripheral nonlinearity on psychometric functions for forward-masked tones. The Journal of the Acoustical Society of America 113:1560–1573.

106. Scheperle RA, Neely ST, Kopun JG, Gorga MP (2008) Influence of in situ, sound-level calibration on distortion-product otoacoustic emission variability. The Journal of the Acoustical Society of America 124:288–300.

107. Schütt HH, Harmeling S, Macke JH, Wichmann FA (2016) Painfree and accurate Bayesian estimation of psychometric functions for (potentially) overdispersed data. Vision Research 122:105–123.

108. Seluakumaran K, Mulders WHAM, Robertson D (2008) Unmasking effects of olivocochlear efferent activation on responses of inferior colliculus neurons. Hearing Research 243:35–46.

109. Shannon RV, Zeng F-G, Kamath V, Wygonski J, Ekelid M (1995) Speech Recognition with Primarily Temporal Cues. Science 270:303–304.

110. Shera CA, Guinan JJ, Oxenham AJ (2002) Revised estimates of human cochlear tuning from otoacoustic and behavioral measurements. Proceedings of the National Academy of Sciences 99:3318–3323.

111. Shera CA, Guinan JJ, Oxenham AJ (2010) Otoacoustic Estimation of Cochlear Tuning: Validation in the Chinchilla. JARO 11:343–365.

112. Skinner PH, Jones HC (1968) Effects of Signal Duration and Rise Time on the Auditory Evoked Potential. Journal of Speech and Hearing Research 11:301–306.

113. Smalt CJ, Heinz MG, Strickland EA (2014) Modeling the Time-Varying and Level-Dependent Effects of the Medial Olivocochlear Reflex in Auditory Nerve Responses. JARO 15:159–173.

114. Smith SB, Cone B (2015) The medial olivocochlear reflex in children during active listening. International Journal of Audiology 54:518–523.

115. Strand JF, Brown VA, Merchant MB, Brown HE, Smith J (2018) Measuring Listening Effort: Convergent Validity, Sensitivity, and Links With Cognitive and Personality Measures. Journal of Speech, Language, and Hearing Research 61:1463–1486.

116. Strickland EA (2001) The relationship between frequency selectivity and overshoot. The Journal of the Acoustical Society of America 109:2062–2073.

117. Strickland EA (2004) The temporal effect with notched-noise maskers: Analysis in terms of input– output functions. The Journal of the Acoustical Society of America 115:2234–2245.

118. Strickland EA (2008) The relationship between precursor level and the temporal effect. The Journal of the Acoustical Society of America 123:946–954.

119. Summerfield Q, Assmann P (1987) Auditory Enhancement in Speech Perception. In: The Psychophysics of Speech Perception (Schouten MEH, ed), pp 140–150. Dordrecht: Springer Netherlands. Available at: 10.1007/978-94-009-3629-4_10 [Accessed July 10, 2024].

120. Swaminathan J, Heinz MG (2012) Psychophysiological Analyses Demonstrate the Importance of Neural Envelope Coding for Speech Perception in Noise. J Neurosci 32:1747–1756.

121. Thibodeau LM (1996) Evaluation of Auditory Enhancement and Auditory Suppression in Listeners With Normal Hearing and Reduced Speech Recognition in Noise. J Speech Lang Hear Res 39:947–956.

122. Tippey KG, Longnecker M (2016) An Ad Hoc Method for Computing Pseudo-Effect Size for Mixed Models. Available at: https://www.semanticscholar.org/paper/An-Ad-Hoc-Method-for-Computing-Pseudo-Effect-Size-Tippey-Longnecker/38521e6da4fe383ff92a01b7abbcabbb440dc462 [Accessed February 15, 2023].

123. Turner CW, Souza PE, Forget LN (1995) Use of temporal envelope cues in speech recognition by normal and hearing-impaired listeners. The Journal of the Acoustical Society of America 97:2568–2576.

124. Viemeister NF (1980) Adaptation of masking. In: Psychophysical, Physiological and Behavioural Studies in Hearing (van den Brink G, Bilsen FA, eds), pp 190–198. Delft University Press.

125. von Klitzing R, Kohlrausch A (1994) Effect of masker level on overshoot in running- and frozen-noise maskers. The Journal of the Acoustical Society of America 95:2192–2201.

126. Walsh KP, Pasanen EG, McFadden D (2010a) Overshoot measured physiologically and psychophysically in the same human ears. Hearing Research 268:22–37.

127. Walsh KP, Pasanen EG, McFadden D (2010b) Overshoot measured physiologically and psychophysically in the same human ears. Hearing Research 268:22–37.

128. Wen B, Wang GI, Dean I, Delgutte B (2012) Time course of dynamic range adaptation in the auditory nerve. Journal of Neurophysiology 108:69–82.

129. Whelan R (2008) Effective Analysis of Reaction Time Data. The Psychological Record 58:475–482.

130. Wichmann FA, Hill NJ (2001a) The psychometric function: I. Fitting, sampling, and goodness of fit. Perception & Psychophysics 63:1293–1313.

131. Wichmann FA, Hill NJ (2001b) The psychometric function: II. Bootstrap-based confidence intervals and sampling. Perception & Psychophysics 63:1314–1329.

132. Willmore BDB, King AJ (2023) Adaptation in auditory processing. Physiological Reviews 103:1025–1058.

133. Winslow RL, Sachs MB (1987) Effect of electrical stimulation of the crossed olivocochlear bundle on auditory nerve response to tones in noise. Journal of Neurophysiology 57:1002–1021.

134. Winslow RL, Sachs MB (1988) Single-tone intensity discrimination based on auditory-nerve rate responses in backgrounds of quiet, noise, and with stimulation of the crossed olivocochlear bundle. Hearing Research 35:165–189.

135. Wojtczak M, Klang AM, Torunsky NT (2019) Exploring the Role of Medial Olivocochlear Efferents on the Detection of Amplitude Modulation for Tones Presented in Noise. JARO 20:395–413.

136. Xu L, Zheng Y (2007) Spectral and temporal cues for phoneme recognition in noise. The Journal of the Acoustical Society of America 122:1758–1764.

137. Xu Y, Cheatham MA, Siegel JH (2017) Identifying the Origin of Effects of Contralateral Noise on Transient Evoked Otoacoustic Emissions in Unanesthetized Mice. Journal of the Association for Research in Otolaryngology 18:543–553.

138. Zhao W, Dhar S (2012) Frequency tuning of the contralateral medial olivocochlear reflex in humans. Journal of Neurophysiology 108:25–30.

139. Zwicker E (1965) Temporal Effects in Simultaneous Masking by White-Noise Bursts. The Journal of the Acoustical Society of America 37:653–663.

